# Projection-defined ventral tegmental area neurons exhibit distinct molecular adaptations during acute fentanyl withdrawal

**DOI:** 10.64898/2026.10.02.756315

**Authors:** Annalisa Montemarano, Hajra Sohail, Meghan E. Hnilo, Logan D. Fox, Arial Briones, Alexandria E. Ostman, Megan E. Fox

**Affiliations:** Department of Anesthesiology and Perioperative Medicine, Pennsylvania State University College of Medicine, Hershey PA; Department of Neuroscience and Experimental Therapeutics, Pennsylvania State University College of Medicine, Hershey PA

**Keywords:** opioid withdrawal, translatomics, gene co-expression network, ventral tegmental area, prefrontal cortex, nucleus accumbens, amygdala

## Abstract

Opioid use disorder (OUD) remains a major public health concern, with potent synthetic opioids driving overdose deaths. Understanding how synthetic opioids like fentanyl alter brain circuits to promote persistent use is critical for advancing OUD treatment. The ventral tegmental area (VTA), which projects to several other brain regions including prefrontal cortex (PFC), nucleus accumbens (NAc) and amygdala (Amy), contributes to both the euphoric and dysphoric phases of OUD. Projection target influences the baseline properties and opioid-induced neuroadaptations of VTA neurons, yet less is known about the molecular adaptations that occur across distinct VTA projections during spontaneous opioid withdrawal. Here, we investigated transcriptional responses to acute spontaneous fentanyl withdrawal in projection-defined VTA neurons. Following chronic fentanyl exposure and 24h of spontaneous withdrawal in male and female mice, we sequenced ribosome-associated RNA from VTA neurons projecting to PFC, NAc, or Amy. We found that fentanyl withdrawal produced divergent transcriptional adaptations in VTA neurons projecting to PFC compared with VTA-NAc and VTA-Amy neurons, whereas adaptations in VTA-NAc and VTA-Amy neurons were more concordant. Weighted gene co-expression network analysis (WGCNA) identified fentanyl-responsive modules across VTA projections, with gene ontology analysis linking these modules to distinct biological processes. We further used regulatory enrichment analysis to identify candidate transcriptional regulators associated with fentanyl-responsive co-expression modules. Together, our data demonstrate that VTA neurons undergo distinct molecular adaptations during fentanyl withdrawal based on their projection target, providing new insight into molecular mechanisms relevant to OUD.

## INTRODUCTION

Opioid use disorder (OUD) is a chronic, relapsing condition characterized by repeated, excessive opioid taking, loss of control over opioid taking, and the emergence of a negative emotional state (dysphoria, anxiety, irritability) during periods of opioid abstinence (Shurman et al., 2010; Koob, 2020). The opioid epidemic in the U.S. has worsened in recent years primarily due to increased prevalence of potent synthetic opioids like fentanyl (Spencer et al., 2024). Fentanyl is pharmacologically unique from other opioids (Eshleman et al., 2020; Sutcliffe et al., 2022; Kelly et al., 2023), and fentanyl exposure is associated with poorer treatment outcomes for OUD (Socias et al., 2022; Kawasaki et al., 2026). Despite this, our understanding of the neural mechanisms of OUD is based largely on natural and semi-synthetic opioids like morphine and heroin, while our understanding of fentanyl-induced neuroadaptations remains limited.

The ventral tegmental area (VTA) is a heterogenous midbrain structure important for mediating both the euphoric and dysphoric phases of OUD (Koob and Volkow, 2010; Koob, 2020). The VTA modulates activity at several other nodes in the reward circuit, sending mostly non-overlapping projections to regions such as the prefrontal cortex (PFC), nucleus accumbens (NAc), and amygdala (Amy). VTA projections to each of these regions are comprised of different proportions of dopaminergic, glutamatergic, GABAergic, and co-transmitting neurons (Taylor et al., 2014; Breton et al., 2019; Phillips et al., 2022; Fox et al., 2024; Fitzgerald and Day, 2025). Projection identity has major influence over VTA neuronal activity, as VTA dopamine neurons vary in their baseline electrophysiological properties (Lammel et al., 2008; Margolis et al., 2008; Chaudhury et al., 2013; Friedman et al., 2014; Juarez and Han, 2016), molecular signatures (Sesack et al., 1998; Lewis et al., 2001; Lammel et al., 2008), and circuit inputs (Beier et al., 2015; Elum et al., 2024) depending on their downstream projection target. VTA dopamine neurons also exhibit projection-specific responses to opioids, including differential degree of recruitment by opioids (Morison et al., 2025), distinct morphine-induced morphological changes (Simmons et al., 2019), and selective inhibition by kappa opioid agonists (Margolis et al., 2006). Importantly, opioid actions in the VTA are not limited to dopamine neurons, as multiple VTA neuronal subtypes express mu opioid receptors (Oriol et al., 2025), participate in opioid seeking behaviors (Bossert et al., 2004; McGovern et al., 2023), and undergo diverse molecular adaptations following opioid self-administration (Browne et al., 2023; Fox et al., 2024). However, it remains unclear whether the projection-specific response to opioids exhibited by VTA dopamine neurons is shared by other VTA neurons, since most characterizations of VTA circuit function focus on dopamine.

Defining VTA neuronal subtypes by characteristics other than neurotransmitter content, such as their circuit output, can help elucidate VTA encoding of behavior. In some cases, projection identity serves as a better predictor of VTA neuronal properties than neurotransmitter identity (Margolis et al., 2008). Furthermore, since opioids engage neuronal ensembles in the VTA that bypass neurotransmitter type (Koutlas et al., 2024; Wang et al., 2024), this approach may be especially useful in the context of opioid-induced neuroadaptations. We previously showed that projection-defined VTA neurons exert differential regulation over fentanyl-context associations (Montemarano et al., 2026). These findings supported distinct roles for VTA neurons that project to PFC (VTA-PFC) or NAc (VTA-NAc) in the motivational aspects of fentanyl misuse, such as fentanyl reinforcement & seeking behavior. However, it remains unclear whether these projection-specific VTA functions extend to hyperkatifeia, or the negative emotional state that emerges from opioid dependence, producing behaviors like anxiety and irritability (Koob and Volkow, 2010; Koob, 2020). VTA projections to amygdala (VTA-Amy) are especially important for hyperkatifeia, as VTA-Amy activity mediates anxiety associated with withdrawal from cocaine (Tian et al., 2022) and alcohol (Klinefelter et al., 2026). The extent to which VTA projections to PFC, NAc, or Amy are differentially implicated in opioid withdrawal remains unknown.

To address these gaps in knowledge, we investigated the molecular adaptations occurring in projection-defined VTA neurons during acute spontaneous withdrawal from chronic fentanyl exposure. We used a chronic fentanyl administration model that produces increased impulsive- and irritability-like behavior following 24h of abstinence. At this timepoint, we used ribosome-associated RNA sequencing to assess translatome changes in VTA neurons projecting to either PFC, NAc, or Amy. We found discordant fentanyl-induced differential gene expression in VTA neurons projecting to PFC compared to those projecting to NAc or Amy. We found substantial diversity in the number and function of differentially expressed genes across the three neuronal populations and identified several projection-specific transcriptional regulators that may contribute to fentanyl-induced gene expression changes. Together, these experiments provide insight into the transcriptional landscape of specific VTA circuits during acute fentanyl withdrawal and identify multiple gene targets with potential implications for fentanyl-induced hyperkatifeia.

## MATERIALS AND METHODS

### Experimental subjects

All procedures were approved by the Institutional Animal Care and Use Committee at the Pennsylvania State University College of Medicine (PSUCOM) and conducted in accordance with NIH guidelines for the use of laboratory animals. C57BL/6J and Rpl22^HA^ (“RiboTag”) mice on a C57BL/6J background were obtained from The Jackson Laboratory (RRID: IMSR_JAX:029977) and bred at PSUCOM. Mice were given food and water *ad libitum* and housed in the PSUCOM vivarium on a 12:12h light:dark cycle with lights on at 7:00 am. All mice were group housed (2-5 mice per cage; same sex and drug group) in corncob bedding and provided with nestlet enrichment. Male and female mice were aged 8-10 weeks at the start of experiments, with sex defined gonadally. Homozygous RiboTag mice were used for RNAseq experiments, and wildtype C57BL/6J mice were used for behavioral experiments.

### Stereotaxic surgery

Adeno-associated virus for Cre recombinase (AAVrg.hSyn.HI.eGFP-Cre.WPRE.SV40) was acquired from Addgene (#105540). AAV (300nL/side) was delivered intracranially to RiboTag mice under isoflurane anesthesia as previously described (Montemarano et al., 2025), using the following coordinates relative to bregma: NAc core (AP +1.6, ML ±1.5, DV −4.4, 10°), medial PFC (AP +1.7, ML ±0.75, DV −2.5, 15°), amygdala ( AP-1.6, ML ±2.7, DV −4.8, 0°). Mice were allowed >2 weeks recovery from stereotaxic surgery before starting i.p. injections.

### Fentanyl injections

Fentanyl citrate (#22659) was obtained from Cayman chemical (Ann Arbor, MI) and was dissolved in sterile saline (0.9%). Mice were weighed on day 1 and received i.p. injections of 2 mg/kg fentanyl, or equivalent volume of saline, once daily for 7 consecutive days. On day 8, mice were weighed again, and percent body weight change was calculated as ([day 8 body weight – day 1 body weight] / day 1 body weight) x100.

### Behavioral testing

On day 8 (∼24-28h after the last i.p. injection), a subset of mice underwent behavioral testing under dim red lighting after acclimating to the testing room. Mice were tested on the elevated zero maze (EZM; Maze Engineers, Skokie, IL), where they were placed in the open arm and recorded for 5 minutes. Any-maze software (Stoelting) was used to automatically quantify time spent in the open and closed arms (apparatus dimensions: 61cm high, 50cm diameter). Following elevated zero maze testing, a subset of mice underwent bottle brush testing (Kimbrough et al., 2017; Sidhu et al., 2018; De Guglielmo et al., 2020; Tortorelli et al., 2026). Mice were placed individually into a clean cage containing a thin layer of bedding, without food, water, or enrichment, and allowed to acclimate for at least 15 min. The wire cage lid was then removed, and mice underwent 10 consecutive bottle- brush trials. For each trial, a bottle brush was advanced from the opposite side of the cage toward the mouse over 1–2 s while being rotated, maintained in contact with the whiskers for 1 s, and then withdrawn over 1–2 s while continuing to rotate. The brush was rotated at the starting position for an additional 1 –2 s and then held stationary for up to 5 s or until the mouse disengaged from the brush. Trials were separated by 10 s. Behavioral responses were scored for investigation of the brush, biting, boxing, following, and tail rattling, which were categorized as aggressive behaviors, as well as escape, digging, jumping, rearing/climbing, defecation, vocalization, and grooming, which were categorized as defensive behaviors.

### RNA isolation & sequencing

On day 8 (24-28h after the last i.p. injection), mice were euthanized by cervical dislocation, and brains were rapidly removed and chilled in ice cold PBS. Cold brains were cut into 1 mm coronal sections using an aluminum brain matrix, and VTA tissue punches were collected with 14-gauge needle, then snap-frozen on dry ice and stored at −80°C until processing. Polyribosomes were immunoprecipitated as described previously (Fox et al., 2020, 2023). Tissue punches from RiboTag mice (3 mice pooled per sample) were homogenized by douncing in 1 mL homogenization buffer, and 50µL supernatant saved as input. Remaining supernatant was incubated with 5 µl anti-HA antibody (BioLegend #901515) at 4°C overnight with constant rotation. Samples were then incubated with 400 µL of protein A/G magnetic beads (Thermo Scientific #88803) overnight at 4°C with constant rotation. Beads were washed in a magnetic rack with high-salt buffer. RNA was extracted using Trizol (Invitrogen) and the RNeasy Mini Kit with a DNAse step (Qiagen) from both immunoprecipitated and input samples by adding lysis buffer supplemented with beta mercaptoethanol and following the manufacturer’s instructions (3-5 samples/sex/projection/drug, 150 total mice). RNA quality and concentration were assessed using an Agilent BioAnalyzer, and cDNA libraries were prepared by the Penn State College of Medicine Genome Sciences Core using the SMARTer Stranded Total RNA-Seq Kit v3 – Pico Input Mammalian (Takara Bio, San Jose, CA) according to the manufacturer’s instructions. Libraries were quantified by qPCR prior to sequencing on an Illumina NovaSeq 6000 with an S2 flow cell, generating 50 base pair, paired-end reads (average of ∼70 million reads per sample). Raw data were demultiplexed using bcl2fastq (Illumina), and adapter trimming was performed after UMI extraction with UMI-tools (Smith et al., 2017). Sequencing reads were aligned to the mouse reference genome (GRCm39, Ensembl release 2.7.1a) using STAR (Dobin et al., 2013) with a maximum of two mismatches allowed per read. Gene-level counts were obtained using featureCounts (Liao et al., 2014), with multi-mapping reads included, reads assigned to multiple overlapping features counted fractionally, and the feature with the largest overlap prioritized.

### Differential gene expression analysis

We applied multiple strategies to characterize differential gene expression. We used limma (lmfit) (Ritchie et al., 2015) in R (v4.5.2) to normalize counts (log2CPM) and determine differential expression between projection neurons vs total VTA with post hoc contrasts (makeContrasts). To assess cell-type enrichment across projections, we used the mean_ratios function from DeconvoBuddies (Huuki-Myers et al., 2025) in conjunction with our previously published snRNAseq dataset of mouse VTA (Fox et al., 2024). The top 50 markers per cluster with a MeanRatio >1 were selected. Overlap between differentially expressed genes and cluster markers was evaluated using Fisher’s exact test. For each projection, the proportion of differentially expressed genes mapping to each marker-defined cell class was calculated as the number of overlapping genes divided by the total number of differentially expressed genes overlapping any cluster marker. We then used lmFit to fit log2 normalized read counts per million to a linear model and tested for effects of fentanyl in each projection target population with sexes combined. To compare differential expression patterns of top DEGs across projections, for each projection we identified 5 DEGs with nominally significant effect (p <0.05) of fentanyl with sexes combined having the highest log fold change, and 5 with the lowest log fold change. To map differential expression patterns of all DEGs across projections, we generated an alluvial plot of all DEGs with nominally significant effect (p <0.05) of fentanyl with sexes combined in R using the easyalluvial package (Koneswarakantha, 2026). To compare differential expression patterns of all genes independent of p value, we conducted threshold free, rank-rank hypergeometric overlap analysis (RRHO) in R using the RRHO2 package (Cahill et al., 2018). Heatmaps were used to visualize regions of significant overlap in gene regulation across projection targets.

### Genome network analysis

To characterize the effects of fentanyl on gene co-expression networks, we used limma to select sets of genes that exhibited nominally significant effect (p <0.05) of fentanyl, sex, or projection target, similar to our previous work (Fox et al., 2023). We then used weighted gene co-expression network analysis (WGCNA) to characterize co-expressed modules, separately within each projection target. Module detection was performed using the blockwiseModules() function with power = 8, corType = bioCor, networkType = signed, minModuleSize = 30, reassignThreshold = 0, mergeCutHeight = 0.25, minMEtoSTay = 0 and otherwise default parameters. We tested for differential expression of module eigengenes with limma, using posthoc contrasts to estimate effects of fentanyl while adjusting for sex differences separately in each target. Hub genes for each module were defined by calculating the Pearson correlation between the eigenene and each gene in the module.

### Gene ontology and transcription factor analysis

We performed Gene Ontology (GO) analysis using gprofiler2 (Kolberg et al., 2020) and queried over-represented GO terms in each WGCNA module using a Benjamini & Hochberg False Discovery Rate (FDR) correction of 0.05. To reduce redundancy and improve biological specificity during visualization, enriched GO terms from g:Profiler analyses were filtered to remove broad parent categories. Terms with large gene sets (term size > 500) were excluded to avoid enrichment driven by highly general processes. For selected WGCNA modules, we used the iRegulon plug-in in Cytoscape (Janky et al., 2014) with default settings to identify candidate transcriptional regulators whose predicted target genes were enriched within each module. To compare candidate transcriptional regulators across projections, we first identified predicted transcription factors that belonged to fentanyl-associated WGCNA modules and were themselves differentially expressed following fentanyl exposure. For each projection we then counted the number of fentanyl-associated target modules each transcription factor was predicted to regulate.

## RESULTS

### Increased impulsive- and irritability-like behavior during acute abstinence from chronic fentanyl

To investigate behavioral and molecular adaptations associated with acute, spontaneous fentanyl withdrawal, we used a chronic fentanyl administration paradigm followed by 24h spontaneous withdrawal. Mice received one week of i.p. fentanyl injections, using a dose previously shown to produce physical dependence, locomotor sensitization, and weight loss (**Fig 1A**) (Uddin et al., 2021). We compared body weight at 24h after the last injection to starting body weight (before the first injection). Consistent with the development of physical dependence, fentanyl-treated mice had lower bodyweight than saline-treated mice (**Fig 1B**, Drug: F_1,57_=6.22, p=0.015). We also assessed somatic withdrawal behavior at the 24-28h timepoint in a pilot cohort; however, escape jumping was observed in only one fentanyl-treated mouse and was therefore not analyzed further.

**Figure 1.**
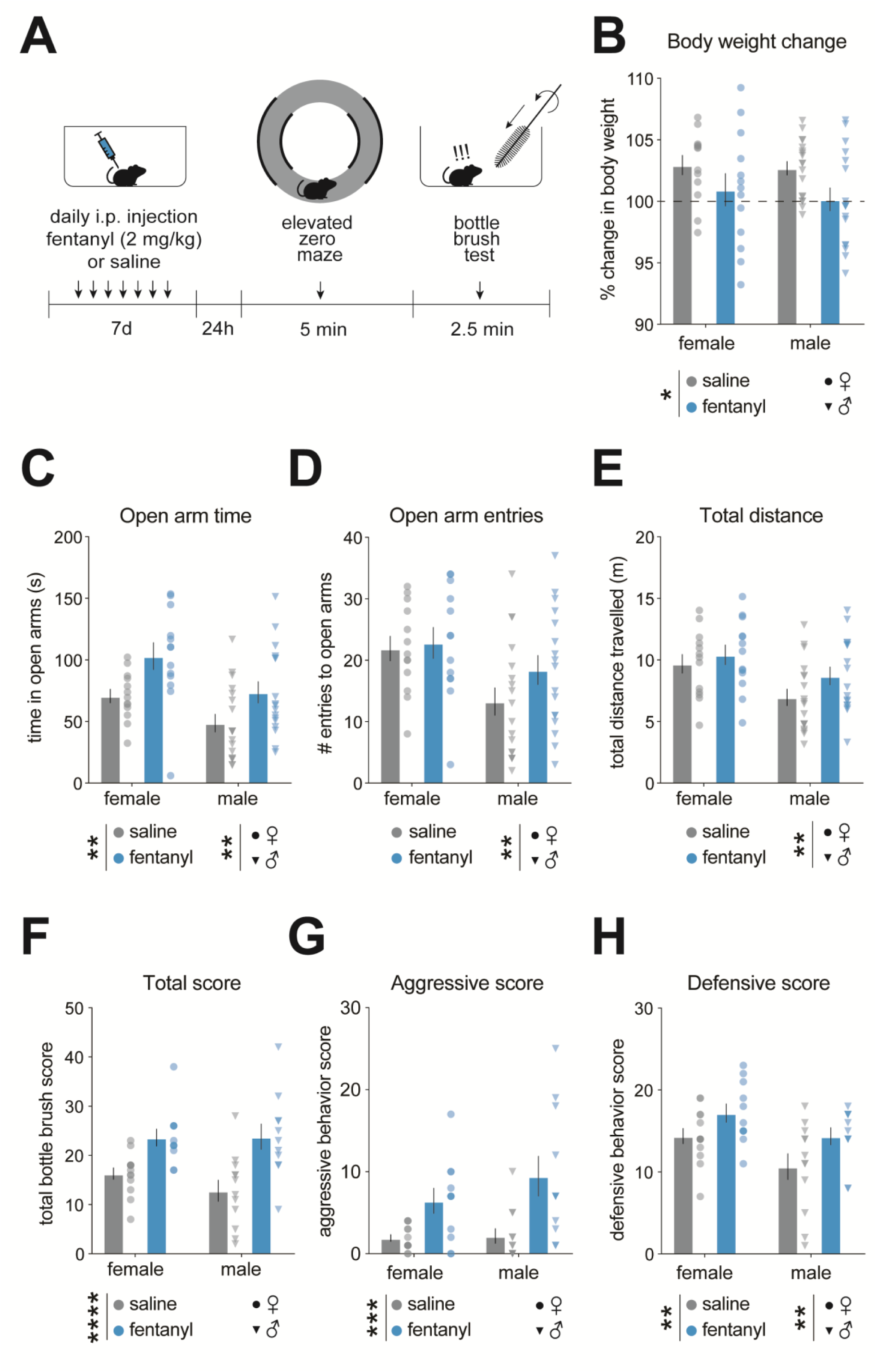
Increased impulsive- and irritability-like behaviors during acute fentanyl withdrawal. **(A)** Experimental timeline. Mice received daily i.p. injections of 2 mg/kg fentanyl or saline for 7 days. On day 8, mice underwent 5 min elevated zero maze (EZM) testing followed >20min later by a 2.5 min bottle brush test. **(B)** Change in body weight change from day 1 (pre-injection) to 8. Fentanyl-treated mice showed lower body weight change than saline-treated mice, which gained weight over the treatment period (n=13-18/sex/treatment, *, p=0.015). **(C)** Fentanyl-treated mice spent more time in the open arms of the EZM than saline-treated mice (**, p=0.0011) and females spent more time in the open arms than males (**, p=0.0033). **(D)** Females made more open-arm entries (**, p=0.0072), and **(E)** traveled a greater total distance (**, p=0.0043) than males, with no significant effects of fentanyl treatment. **(F)** In the bottle brush test, fentanyl-treated mice had higher total behavioral response scores (n=11-13/sex/treatment, ****, p<0.0001), **(G)** aggressive behavior scores (***, p=0.0002), and **(H)** defensive behavior scores (**, p=0.0095) than saline-treated mice. Females also had higher defensive scores than males (**, p=0.0091). Data are mean±SEM with individual mice represented as circles (females) or triangles (males). Stars denote main effect of drug or sex; no significant drug x sex interactions were detected.

To assess behavioral responses during acute spontaneous abstinence, we tested mice on the elevated zero maze (EZM) 24-28h after the last fentanyl injection. We found fentanyl-treated mice spent more time in the open arms of the EZM than saline mice, and also that females spent more time in the open arms than males (**Fig 1C**, Drug: F_1,57_=11.73, p=0.0011, Sex: F_1,57_=9.432, p=0.0033). Females also made more open-arm entries (**Fig 1D**, Sex: F_1,57_=7.76, p=0.0072) and travelled a greater overall distance on the EZM (**Fig 1E**, Sex: F_1,57_=8.84, p=0.0043) than males, with no significant drug effects on either measure.

Increased open arm time is typically interpreted as either decreased anxiety-like behavior or increased behavioral disinhibition. To determine whether our findings were more consistent with behavioral disinhibition, we next assessed irritability using the bottle brush test with a subset of mice. Behavioral responses in the bottle brush test are categorized into “aggressive” or “defensive,” with total irritability expressed as the sum of all behavioral responses. First, we found fentanyl mice had greater overall irritability score compared to saline mice (**Fig 1F**, Drug: F_1,43_=22.35, p<0.0001). When breaking this down by behavior type, we found fentanyl mice exhibited increased aggressive (**Fig 1G**, **Fig 1F**, Drug: F_1,43_=17.05, p=0.0002) and defensive (**Fig 1H**, Drug: F_1,43_=7.36, p=0.0095) responses. There were sex differences in defensive score, with females exhibiting more defensive responses than males (**Fig 1H**, Sex: F_1,43_=7.46, p=0.0091).

### Discordant fentanyl-induced transcriptional adaptations in projection-specific VTA neurons

VTA projection neurons exhibit distinct morphologic adaptations and differential immediate early gene induction following chronic opioid exposure (Simmons et al., 2019; Morison et al., 2025). To examine if similar projection-specific adaptations occur at the transcriptional level, we used a retrograde Cre virus in either PFC, NAc, or Amy in Ribotag mice to express HA tagged ribosomes in specific VTA projection neurons. Mice received one week of fentanyl injections as above. After 24-28h abstinence, we isolated and sequenced ribosome-associated RNA from VTA neurons projecting to either PFC, NAc, or Amy, along with RNA from the input fraction representing total VTA (**Fig 2A**). First, to confirm projection neuron enrichment, we compared expression of several cell-type markers in the immunoprecipitated samples relative to input in saline-treated mice. We found significant enrichment of general neuronal markers (*Mapt*, *Tubb3*, *Map2*, *Nlgn3*, *Nefl, Eno2,* p’s≤ 1.1×10^-8^), and de-enrichment of markers for oligodendrotyces (*Aspa*), astrocytes (*Slc1a3*), microglia (*Arhgap15*), and pericytes (*Pla1a*, p’s≤ 2.9×10^-4^, **Supplemental Fig 1A** and **Supplemental Files.**). We next looked for enrichment of neuron subtype markers by comparing with mouse VTA neuron clusters previously identified by our group (Fox et al., 2024). Across all 3 projection populations, we found significant enrichment for marker genes associated with dopamine, glutamate, and GABA neuron clusters (**Supplemental Fig 1B** and **Supplemental Files**). Most neuron cluster marker sets were similarly enriched across projections, although some showed projection-specific patterns. Markers associated with the co-transmission/combinatorial dopamine cluster (DA-combo-18) were enriched in VTA-PFC and VTA-NAc but were not significantly enriched in VTA-Amy. One glutamate cluster (GLUT-11) was uniquely enriched in VTA-PFC, whereas two glutamate clusters (GLUT-20 and GLUT-26) were enriched in VTA-NAc and VTA-Amy but not VTA-PFC. Finally, markers associated with one GABA cluster (GABA- 24) were uniquely enriched in VTA-Amy.

**Figure 2.**
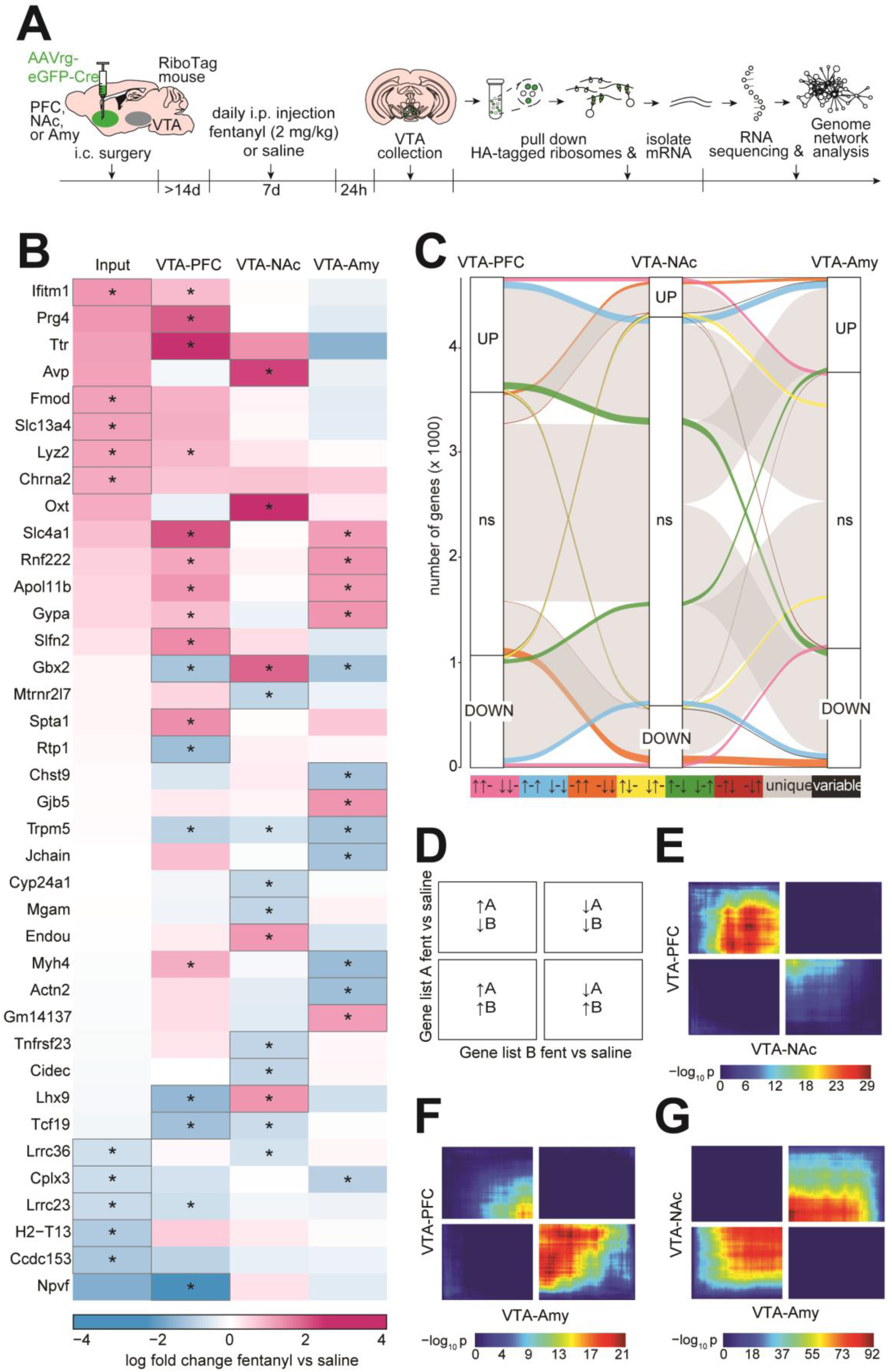
Fentanyl withdrawal induces discordant transcriptional adaptations among VTA projection neurons. **(A)** Timeline for projection-specific VTA RNA sequencing experiments. RiboTag mice received retrograde Cre in either medial PFC, NAc core, or amygdala (n=3-5 samples/sex/drug/projection group; 3 mice pooled per sample; total mice: VTA-PFC, n=51; VTA-NAc, n=51; VTA-Amy n=48). Two weeks later, mice received daily i.p. injections of 2 mg/kg fentanyl or equivalent volume of saline for 7 days. Approximately 24-28 hours after the last injection, brains were collected and fresh tissue punches containing VTA were flash frozen. HA-tagged ribosomes from VTA projection neuron populations were immunoprecipitated and purified RNA was used to prepare cDNA libraries for RNA sequencing. **(B)** Heatmaps showing top 5 upregulated and top 5 downregulated DEGs (gray outline) in fentanyl vs saline (*, p<0.05) across Input (total VTA), VTA-PFC, VTA- NAc, and VTA-Amy samples. **(C)** Alluvial plot comparing DEGs between neuronal populations. Combinations of upregulated (↑), downregulated (↓), and not significant (-) genes are represented by individual colors. DEGs in the same direction across two projections and not significant in a third are represented in pink, blue, and orange. DEGs in opposite directions across two projections and not significant in a third are represented in yellow, green, and red. DEGs that are uniquely up or down in a single population are represented in gray, and the remaining combinations are represented in black. **(D)** RRHO interpretation schematic. **(E)** RRHO plots showing discordance of fentanyl-induced DEGs between VTA-PFC and VTA-NAc, **(F)** discordance between VTA-PFC and VTA-Amy, and **(G)** concordance between VTA-NAc and VTA-Amy, with −log_10_(P value) encoded in color using the adjacent color bars.

We next examined the top differentially expressed protein coding genes in each VTA projection population following fentanyl exposure and compared them with the top differentially expressed genes (DEGs) in total VTA (**Fig 2B**, all genes in **Supplemental Files**). There was very little overlap in the top 5 most up- and downregulated genes among the projections. For example, *Gbx2* and *Lhx9*, respectively encoding the transcription factors gastrulation brain homeobox 2 and LIM homeobox 9, were two of the top upregulated genes in VTA-NAc, but were among the most *downregulated* in VTA-PFC. Several genes on the list were similarly upregulated in both VTA-PFC and VTA-Amy, including *Slc4a1* (encoding anion exchanger 1), *Rnf222* (encoding ring finger protein 222), *Apol11b* (encoding apolipoprotein L 11b), and *Gypa* (encoding glycophorin A). The only gene on this list that was similarly changed across all 3 projection populations was *Trpm5*, encoding a member of the transient receptor potential cation channel family. *Trpm5* was among the top downregulated genes in VTA-Amy and was also significantly downregulated in both VTA-PFC and VTA-NAc. When extending the comparison to *any* DEG with p<0.05, there was still very little overlap among the projection populations, with most genes being uniquely up- or downregulated in a single projection (**Fig 2C**). With this nominal cutoff, nearly twice as many genes were differentially expressed in VTA-PFC and VTA-Amy (VTA-PFC: 1091 upregulated, 1065 downregulated; VTA- Amy: 900 upregulated, 1132 downregulated) compared to VTA-NAc (377 upregulated, 587 downregulated; **Fig 2C & Supplemental Files**). There was a comparable number of DEGs that changed in the same or opposite directions when comparing VTA-PFC to VTA-NAc and VTA-Amy. However, comparison of VTA-NAc to VTA-Amy revealed a greater number of DEGs that changed in the same direction than in opposite directions (**Fig 2C**).

To ensure these observations were not driven by arbitrary p-value cutoffs, we next applied rank-rank hypergeometric overlap (RRHO) analysis. This threshold-free method assesses the significance of overlap between two gene lists ranked by their degree of differential expression rather than by p-value (Plaisier et al., 2010; Cahill et al., 2018). We first compared fentanyl-induced gene expression changes in input to each projection population and expected significant overlap as these populations are part of total VTA. For all 3 comparisons we found significant overlap in the upper right and lower left quadrants, indicating concordant patterns of differential expression (**Supplemental Fig 1C-E**). When comparing ranked gene lists between VTA-PFC and VTA-NAc, we found robust overlap in the upper left and lower right quadrants, revealing discordant patterns of differential expression as a function of fentanyl (**Fig 2E**). We also found discordance when comparing fentanyl DEGs between VTA-PFC and VTA-Amy (**Fig 2F**). In contrast, fentanyl DEGs within VTA-NAc and VTA- Amy were concordant, reflected by overlap in upper right and lower left quadrants of the RRHO plot, and consistent with our analysis using p<0.05 cutoff (**Fig 2G**). Taken together, these data suggest that while each projection population undergoes unique transcriptional adaptations, changes in VTA-NAc and VTA-Amy are more similar to one another than to VTA-PFC.

### Fentanyl differentially alters the molecular landscape of VTA neurons depending on downstream projection target

To further characterize transcriptional adaptations in VTA projection neurons, we next performed weighted gene co-expression network analysis (WGCNA) separately in each projection neuron population. We identified 23 modules of co-expressed genes in VTA-PFC, 12 in VTA-NAc, and 20 in VTA-Amy that showed a significant (p<0.05) effect of fentanyl in either sex analyzed separately or with sexes combined (**Fig 3A-C, Supplemental Files**). To define the functional enrichment within the fentanyl associated modules, we performed gene ontology (GO) analysis on the genes within each module (Select terms in **Fig 3D-F**, all enrichments in **Supplemental Files**). Genes in upregulated VTA-PFC modules were enriched for GO terms relating to intracellular signal transduction and structural remodeling, while downregulated VTA-PFC modules included terms relating to synaptic signaling, calcium regulation, intracellular trafficking, and cellular stress response (**Fig 3D**). While there were fewer enriched GO terms for VTA-NAc, their functions broadly reflected an upregulation of transcription, and a downregulation of intracellular metabolism and protein trafficking (**Fig 3E)**. Lastly, VTA- Amy upregulated modules were enriched for GO terms relating to neurotransmitter transport and energy metabolism, and downregulated modules enriched for protein degradation and genome maintenance ( **Fig 3F**).

**Figure 3.**
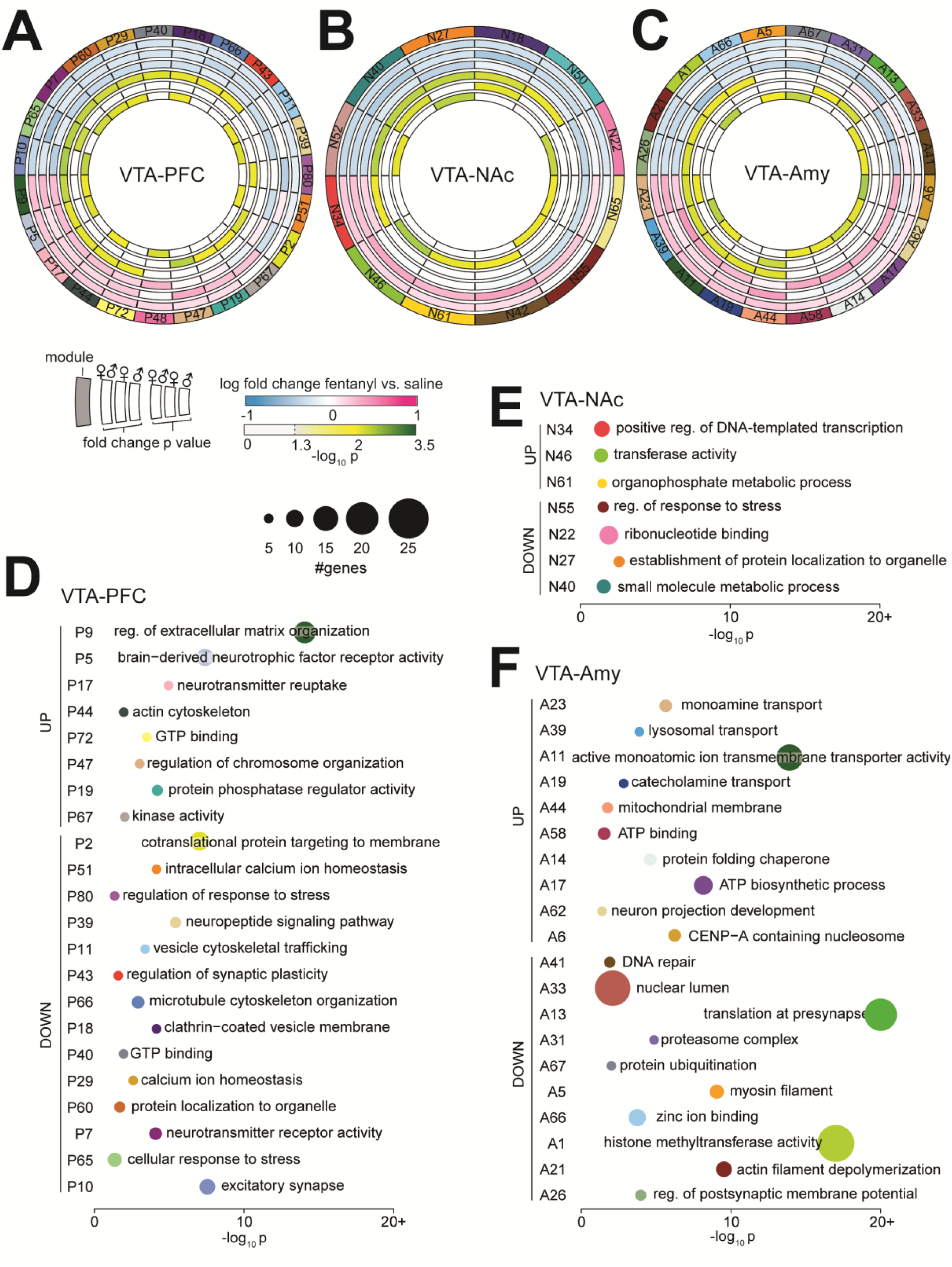
Fentanyl withdrawal-induced DEG function varies among VTA projections. **(A)** Circos plot showing fentanyl-associated WGCNA modules in VTA-PFC, **(B)** VTA-NAc and **(C)** VTA-Amy, defined as differential eigengene expression with p<0.05 effect of fentanyl, ranked clockwise by log fold change. Outer ring indicates an individual module assigned an arbitrary color and number. Fold change in eigengene expression and −log_10_(p value) are shown internally as sexes combined, then separately in females, and males, respectively. Hashed line on color bar indicates p<0.05 cut-offs. **(D)** Gene ontology analysis of VTA-PFC fentanyl modules. Modules are sorted by the most upregulated (UP) to downregulated (DOWN). Circle size represents number of genes in the module enriched for the GO term. One significant GO term was chosen to represent each module. **(E)** Gene ontology analysis of VTA-NAc fentanyl modules. **(F)** Gene ontology analysis of VTA-Amy fentanyl modules.

We next visualized four gene networks per projection population, based on the top upregulated and downregulated eigengenes, along with an upregulated and downregulated module enriched for GO terms of interest. For VTA-PFC, the top upregulated fentanyl-associated module P9—which was enriched for GO term “regulation of extracellular matrix organization”—contained hub genes encoding nidogen-1 (*Nid1*) and fibronectin-1 (*Fn1* **Fig 4A**). An additional upregulated module, P5, enriched for GO term “brain-derived neurotrophic factor receptor activity” contained a hub gene encoding the transcriptional repressor Rev-Erbβ (*Nr1d2,* **Fig 4B**). The most downregulated VTA-PFC module, P10, was enriched for the GO term “excitatory synapse” and contained hub genes encoding calretinin (*Calb2*) and synaptotagmin-13 (*Syt13*, **Fig 4C**). An additional downregulated VTA-PFC module, P39—which was enriched for “neuropeptide signaling pathway”— contained a hub gene encoding the galanin receptor-1 (*Galr1* **Fig 4D**). Since transcription factor-encoding genes were among the most discordant DEGs between projection populations (**Fig 2B**), we identified potential upstream transcriptional regulators of our select gene modules with iRegulon (Janky et al., 2014). We chose to visualize transcription factors that belonged to the module of interest and were also differentially expressed as a function of fentanyl exposure (select transcription factors in **Fig 4E, 4F**, all in **Supplemental Files**). For VTA-PFC we found *Nfkb2* was significantly upregulated in P5 and a predicted regulator of the genes in P5 (**Fig 4E**). Similarly, *Barx2* was significantly downregulated in P39 and is a predicted regulator of P39 (**Fig 4F**).

**Figure 4.**
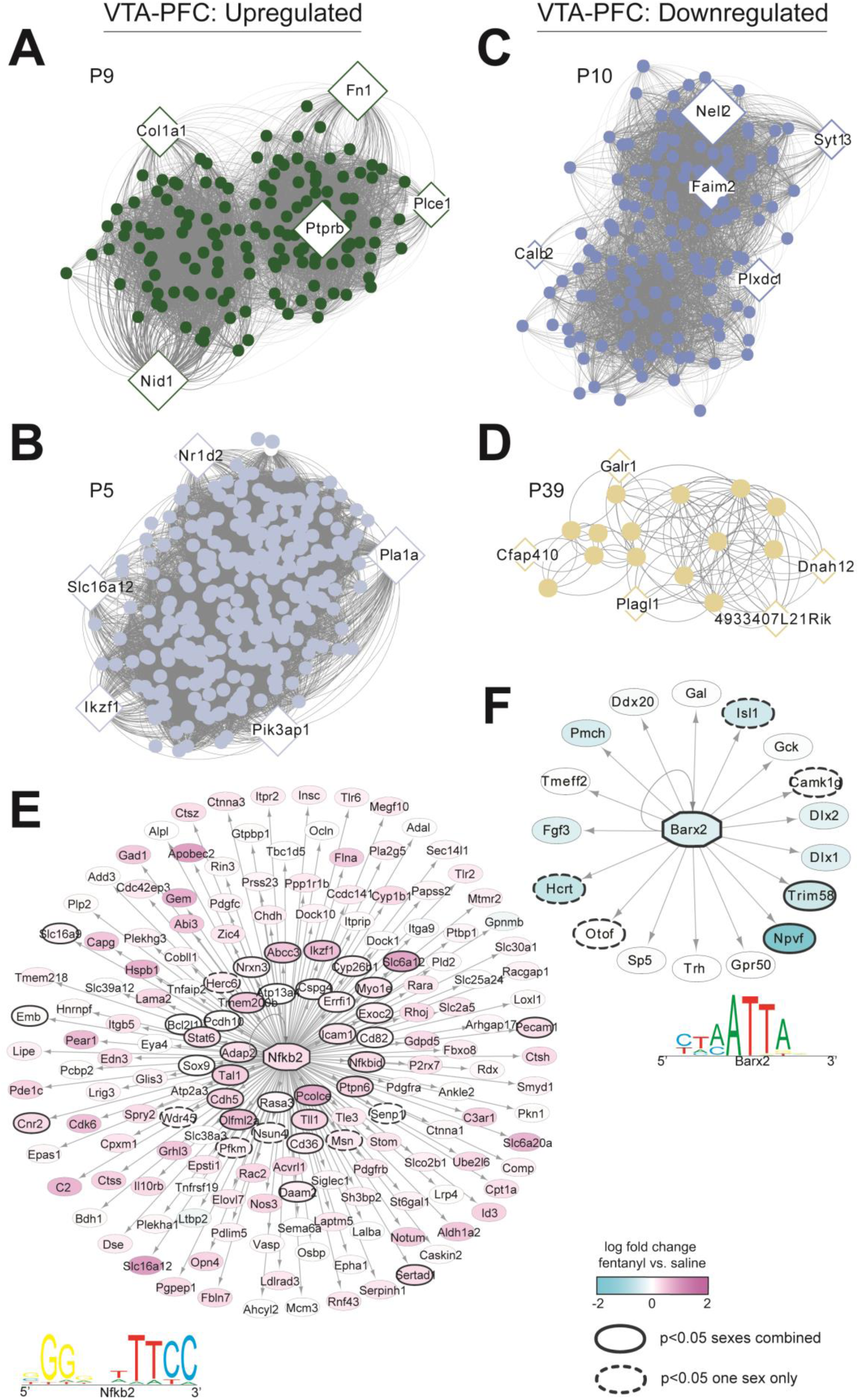
VTA-PFC gene networks and transcriptional regulators altered by fentanyl withdrawal. **(A)** Gene network for P9, the VTA-PFC fentanyl-associated module with the largest positive module eigengene fold change. Genes with the most connections (hub genes) are labeled and represented by diamonds, with diamond size relatively scaled to the number of connections. **(B)** Gene network for P5, an additional upregulated VTA- PFC fentanyl-associated module. **(C)** Gene network for P10, the VTA-PFC fentanyl-associated module with the largest negative module eigengene fold change. **(D)** Gene network for P39, an additional downregulated VTA- PFC fentanyl-associated module. **(E)** Regulatory network for *Nfkb2*, a candidate transcriptional regulator of P5. Arrows indicate P5 genes predicted to be regulatory targets of *Nfkb2*. Color represents the log fold change of each gene following fentanyl exposure. Solid black outline indicates genes significantly altered by fentanyl (p<0.05) when sexes were analyzed together and hashed black outlines indicate genes significantly altered in one sex only. A representative enriched binding motif for *Nfkb2* is shown below. **(F)** Regulatory network for *Barx2*, a candidate regulator of P39, with differentially expressed genes denoted as in (E). *Barx2* representative binding motif is shown below.

For VTA-NAc, the most upregulated fentanyl-associated module, N34, enriched for the GO term “positive regulation of DNA-templated transcription” contained hub genes encoding multiple transcriptional regulators (*Zfp207*, *Cdk5rap3*, *Bacc1*, **Fig 5A**). An additional upregulated module, N46, enriched for GO term “transferase activity,” had hub genes involved in nuclear transport (*Nup35, Hcfc1r,* **Fig 5B**) and potassium flux (*Kcnk4*, **Fig 5B**). The most downregulated VTA-NAc fentanyl module, N52, was not significantly enriched for a GO term. Instead, we visualized the network for the second (N40, “small molecule metabolic process”) and third (N27, “establishment of protein localization to organelle”) most downregulated VTA-NAc modules. These networks contained hub genes encoding transcriptional regulators (*Zfp296*, **Fig 5C**; *Cdk9* **Fig 5D**), as well as proteins involved in DNA repair (*Zswim7*, **Fig 5C**), mRNA transport (*Gle1,* **Fig 5D**), calcium sensing (*Calm3*, **Fig 5D**), and proteostasis (*Fbxw5*, **Fig 5C**; *Dnajb9,* **Fig 5D**). When identifying DEGs that are predicted transcriptional regulators of genes in VTA-NAc modules, we found *Egr1*, which was significantly upregulated in N34 and a predicted regulator of genes in N46 (**Fig 5E**), as well as *Gabpb1*, which was significantly downregulated in N18 and a predicted regulator of genes in N55 (**Fig 5F**).

**Figure 5.**
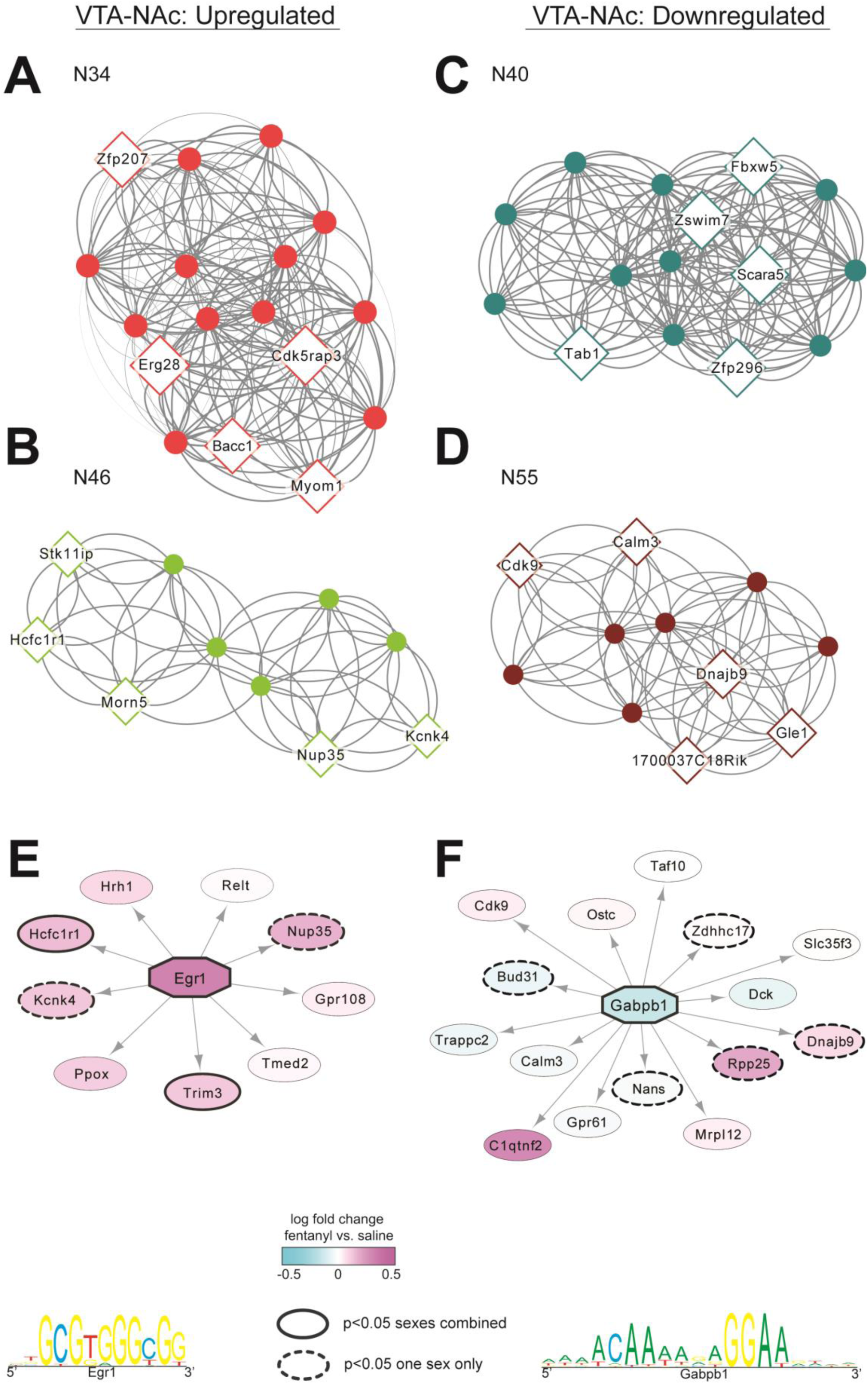
VTA-NAc gene networks and transcriptional regulators altered by fentanyl withdrawal. **(A)** Gene network for N34, the VTA-NAc fentanyl-associated module with the largest positive module eigengene fold change. Genes with the most connections (hub genes) are labelled and represented by diamonds, with diamond size relatively scaled to the number of connections. **(B)** Gene network for N46, an additional upregulated VTA-NAc fentanyl-associated module. **(C)** Gene network for N40, the VTA-NAc fentanyl- associated module with the largest negative module eigengene fold change. **(D)** Gene network for N55, an additional downregulated VTA-NAc fentanyl-associated module. **(E)** Regulatory network for *Egr1*, a candidate transcriptional regulator of N46. Arrows indicate N46 genes predicted to be regulatory targets of *Egr1*. Color represents the log fold change of each gene following fentanyl exposure. Solid black outline indicates genes significantly altered by fentanyl (p<0.05) when sexes were analyzed together and hashed black outlines indicate genes significantly altered in one sex only. A representative enriched binding motif for *Egr1* is shown below. **(F)** Regulatory network for *Gabpb1*, a candidate regulator of N55, with differentially expressed genes denoted as in (E). *Gabpb1* representative binding motif is shown below.

For VTA-Amy, the most upregulated module was A23 and enriched for the GO term “monoamine transport.” Accordingly, the A23 network contained hub genes encoding DOPA decarboxylase (*Ddc*), vesicular monoamine transporter 2 (*Slc18a2*), 5-HT receptor 1A (*Htr1a*, **Fig 6A**). An additional upregulated module, A19, was enriched for “catecholamine transport” and contained a hub gene encoding the equilibrative nucleoside transporter 2 (*Slc29a2*, **Fig 6B**). The gene for the equilibrative nucleoside transporter 1, *Slc29a1*, appeared as a hub gene in the most downregulated VTA-Amy network, A26 (**Fig 6C**). A26 was enriched for “regulation of postsynaptic membrane potential,” and the second most downregulated VTA-Amy network, A21, was enriched for “actin filament depolymerization.” These downregulated VTA-Amy modules contained hub genes encoding proteins involved in G-protein signaling (*Rgs16,* **Fig 6C**), DNA repair (*Elof1*, **Fig 6C**), RNA processing (*Nxf1*, **Fig 6C**; *Ewsr1*, **Fig 6D**), and ubiquitination (*Usp43*, **Fig 6C**; *Marchf7*, **Fig 6D**). When identifying VTA-Amy DEGs that may transcriptionally regulate genes in VTA-Amy modules, we found *Nhlh1* (**Fig 6E**) and *Neurod1* (**Fig 6F**), which were respectively upregulated in A23 and downregulated in A26, and predicted regulators of their respective modules.

**Figure 6.**
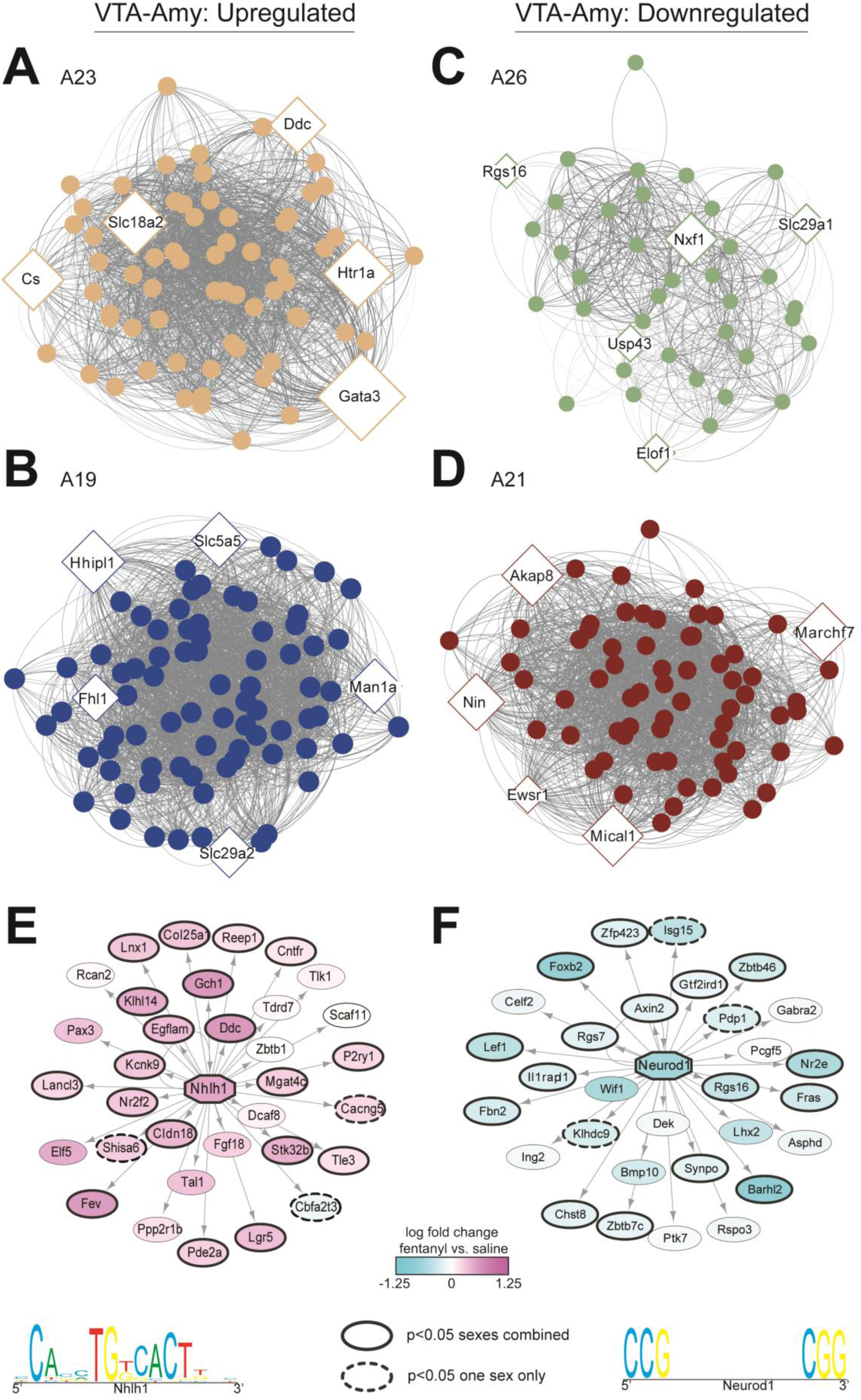
VTA-Amy gene networks and transcriptional regulators altered by fentanyl withdrawal. **(A)** Gene network for A23, the VTA-Amy fentanyl-associated module with the largest positive module eigengene fold change. Genes with the most connections (hub genes) are labelled and represented by diamonds, with diamond size relatively scaled to the number of connections. **(B)** Gene network A19, an additional upregulated VTA-Amy fentanyl-associated module. **(C)** Gene network for A26, the VTA-Amy fentanyl-associated module with the largest negative module eigengene fold change. **(D)** Gene network for A21, an additional downregulated VTA-Amy fentanyl-associated module. **(E)** Regulatory network for *Nhlh1*, a candidate transcriptional regulator of A23. Arrows indicate A23 genes predicted to be regulatory targets of *Nhlh1*. Color represents the log fold change of each gene following fentanyl exposure. Solid black outline indicates genes significantly altered by fentanyl (p<0.05) when sexes were analyzed together and hashed black outlines indicate genes significantly altered in one sex only. A representative enriched binding motif for *Nhlh1* is shown below. **(F)** Regulatory network for *Neurod1*, a candidate regulator of A26, with differentially expressed genes denoted as in (E). *Neurod1* representative binding motif is shown below.

Given the abundance of DEGs in our dataset that were predicted transcriptional regulators of other DEGs, we next wanted to determine how fentanyl-altered transcriptional regulation varied across VTA projection populations. For all genes that were significantly upregulated (**Fig 7A**) or downregulated (**Fig 7B**) in any fentanyl-associated module, we identified the number of fentanyl modules they were predicted to regulate (target fentanyl-associated modules). While both VTA-PFC and VTA-Amy contained differential expression of many predicted transcriptional regulators, they each regulate only a few (1-3) target fentanyl modules each. In contrast, while VTA-NAc contained fewer DEGs encoding transcription factors, they were predicted to regulate many (up to 7- 8) target VTA-NAc fentanyl modules. Further, while VTA-PFC and VTA-Amy had multiple shared predicted regulators (*Gata3*, *Barhl2*, *Nkx1-1*), VTA-NAc transcriptional regulators did not overlap with any other projections.

**Figure 7.**
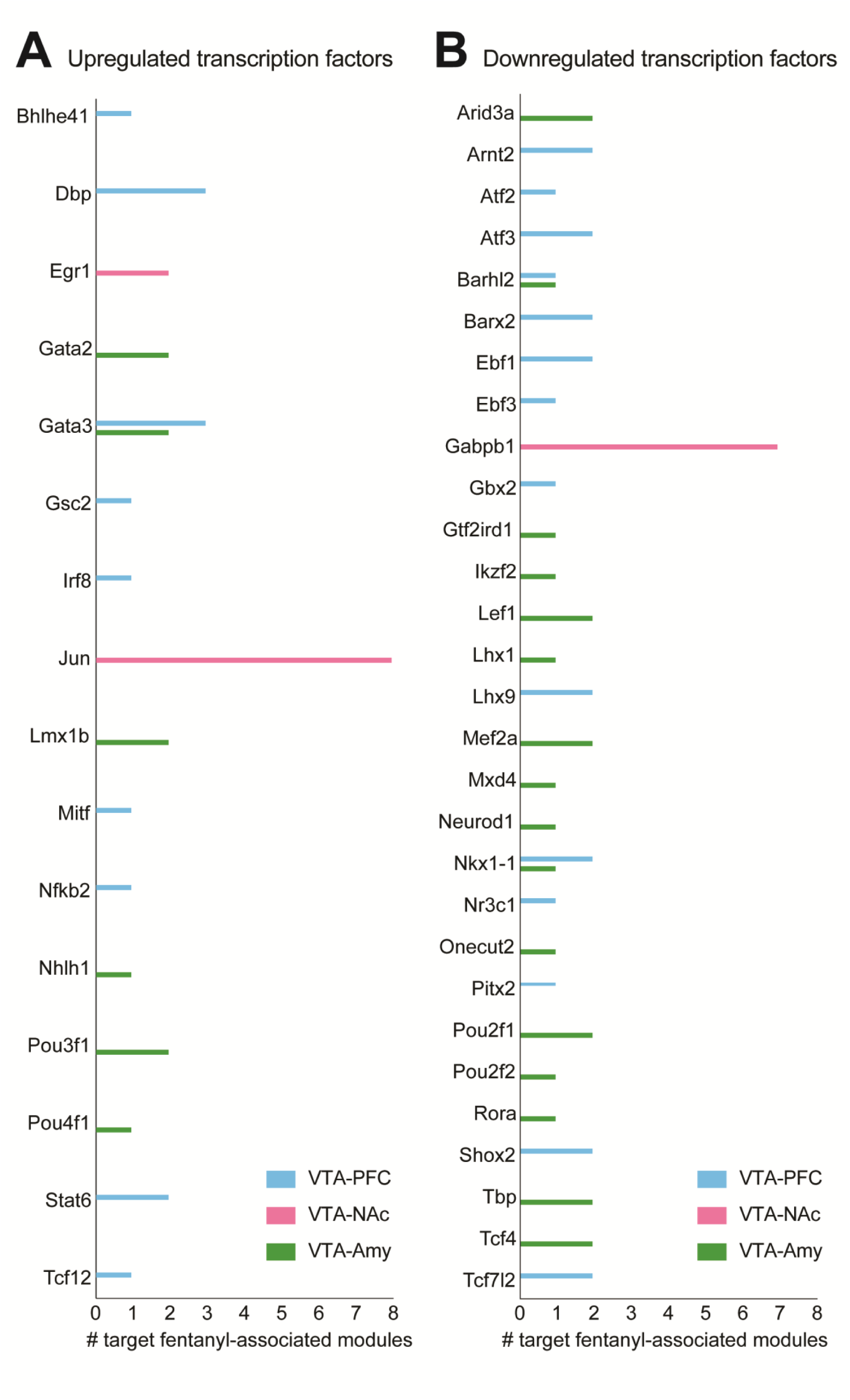
Projection-specific patterns of candidate transcriptional regulation during fentanyl withdrawal. **(A)** Upregulated differentially expressed genes (p<0.05 fentanyl effect) identified as candidate transcriptional regulators of fentanyl-responsive modules. Bars indicate the number of modules associated with each candidate regulator in VTA-PFC (blue), VTA-NAc (pink), and VTA-Amy (green). **(B)** Downregulated differentially expressed genes (p<0.05 fentanyl effect) identified as candidate transcriptional regulators of fentanyl-responsive modules, displayed as in (A).

## DISCUSSION

In this study we demonstrated that acute, spontaneous fentanyl withdrawal induces distinct molecular adaptations in VTA neurons depending on their downstream projection target. We used RNA sequencing to show that fentanyl induces discordant translatomes in VTA neurons that project to the cortex (VTA-PFC) vs those that project to limbic regions (VTA-NAc and VTA-Amy). We found that fentanyl gene modules are associated with a variety of cellular functions that differ among VTA projection populations. Further, we identified multiple transcription factors with implications for fentanyl-induced transcriptional adaptations. These experiments emphasize how the transcriptional landscape of the VTA vastly differs based on circuit output and lay the foundation for future mechanistic studies of how molecular adaptations across VTA networks contribute to fentanyl withdrawal.

### Impulsivity & irritability during acute fentanyl withdrawal

To assess behavioral responses associated with acute, spontaneous withdrawal from chronic fentanyl exposure, we performed EZM and bottle brush tests ∼24h after the last fentanyl injection. We found mice that had received fentanyl had increased time in the open arms of the EZM relative to saline mice, suggesting an anxiolytic effect. This finding was somewhat unexpected, since acute morphine withdrawal classically produces increased anxiety-like behavior, reflected by reduced open-arm time in rats (Bhattacharya et al., 1995; Schulteis, 1998), although the opposite is true during protracted morphine withdrawal in mice (Bravo et al., 2020), Alternatively, increased exploration of the open arms during opioid withdrawal can instead be interpreted as an increased impulsivity or risk-taking behavior (Bravo et al., 2020). Indeed, rats undergoing prolonged forced abstinence from fentanyl self-administration demonstrate increased risky behavior (Wheeler et al., 2023; Hrelja et al., 2025). Given this, our EZM findings suggest similar impulsivity or risk-taking behavior may be emerging earlier in the time course of fentanyl abstinence, though the specific time course of anxiety-like behavioral expression through spontaneous fentanyl withdrawal has not been fully investigated. We also found fentanyl-treated mice performed more aggressive & defensive behavioral responses in the bottle brush test compared to saline-treated mice, suggesting increased irritability-like behavior. Increased irritability in the bottle brush test is also seen during withdrawal from chronic alcohol (Sidhu et al., 2018) and oxycodone (De Guglielmo et al., 2020), and hyperirritability is a classic somatic withdrawal sign in rats undergoing precipitated withdrawal from morphine (Schulteis et al., 1997) and fentanyl (Uddin et al., 2021). While somatic withdrawal signs are readily observed in animals undergoing naloxone-precipitated opioid withdrawal, they were infrequent at the 24-28h spontaneous fentanyl withdrawal timepoint assessed here, leading us to prioritize other behavioral measures. Future work, especially those incorporating machine-learning based behavioral analysis, will be important for defining the time course and spectrum of somatic withdrawal signs following fentanyl exposure across species, given its distinct pharmacokinetic properties and high lipophilicity relative to natural opioids.

### Projection-specific VTA neurons undergo discordant fentanyl-induced molecular adaptations

To investigate fentanyl-induced molecular adaptations in projection-defined VTA neurons, we sequenced ribosome-associated RNA from VTA neurons that project to one of three major target regions, either PFC, NAc, or Amy, after chronic fentanyl exposure. We found surprisingly little overlap when comparing the top 5 most upregulated and most downregulated DEGs among these neuronal populations. The only two shared genes on this list, the transcription factor-encoding genes *Gbx2* and *Lhx9*, were two of the most upregulated DEGs in VTA-NAc and two of the most downregulated DEGs in VTA-PFC. Interestingly, the only gene on this list that was similarly altered among all 3 projections was *Trpm5*, which was one of the most downregulated VTA-Amy genes and was also significantly downregulated in VTA-PFC and VTA-NAc. *Trpm5* encodes a transient receptor potential (TRP) melastatin channel, and while TRP channel activity in several brain regions including PFC, NAc, and Amy has been implicated in substance use disorders (Wescott et al., 2013), their role in the VTA is less studied and warrants further investigation. Our analysis also revealed several DEGs with prior implications for substance use disorder. Some examples include *Slc13a4* and *Ttr*, which are upregulated in input and VTA-PFC respectively in our dataset and are also upregulated in other brain regions after cocaine exposure (Xu et al., 2012; Feng et al., 2014; Navandar et al., 2021). Others, such as *Tcf19* and *Cplx3,* which were downregulated in multiple VTA populations in our dataset are also downregulated in PFC after cocaine exposure (Winstanley et al., 2007; Nezhyva et al., 2024). Since these genes are similarly altered across multiple brain regions and with multiple substances, they make promising molecular targets for future substance use disorder research.

Beyond just the top DEGs, we further compared all significant (p<0.05) DEGs, revealing additional patterns of differential expression. Comparing VTA-NAc to VTA-Amy revealed a greater number of DEGs that changed in the same direction than in opposite directions. This was in line with the threshold-free RRHO analysis, which revealed concordant patterns of differential expression between VTA-NAc and VTA-Amy. VTA-PFC, on the other hand, exhibited discordant patterns of differential expression when compared VTA-NAc and VTA-Amy. Taken together, these findings suggest that during fentanyl withdrawal, VTA neurons projecting to NAc or Amy undergo more similar transcriptional adaptations to each other than compared to PFC-projecting neurons. It is perhaps unsurprising that VTA neurons projecting the cortex are largely different than those projecting to limbic regions, as midbrain dopamine neurons projecting to PFC have unique electrophysiological and molecular properties compared to those projecting to NAc or Amy, while NAc- and Amy-projecting dopamine neurons exhibit more similar properties (Lammel et al., 2008; Margolis et al., 2008). Our findings here suggest these patterns also extend to fentanyl-induced molecular changes.

### Projection-specific gene expression changes reveal distinct cellular functions associated with fentanyl withdrawal

A notable difference between VTA projections includes the biological processes associated with genes altered during fentanyl withdrawal. In VTA-PFC, GO analysis identified enrichment for terms related to neuronal plasticity and excitability, suggesting that fentanyl-responsive genes in this projection may participate in these processes. Similar opioid-induced adaptations occur in VTA dopamine neurons (Doyle and Mazei-Robison, 2021), but the extent to which they differ across projection-defined populations remains unclear. Terms like “regulation of extracellular matrix organization” and “actin cytoskeleton” were enriched across several upregulated VTA-PFC modules, raising the possibility that these genes contribute to structural plasticity. This is reminiscent of chronic morphine increasing soma size exclusively in VTA-PFC dopamine neurons (Simmons et al., 2019) and increasing dendrite diameter of dopamine neurons in the parabrachial VTA, where PFC-projecting neurons primarily reside (Lane et al., 2008). Conversely, several downregulated VTA-PFC modules were enriched for terms related to calcium homeostasis and synaptic function (“intracellular calcium ion homeostasis,” “calcium ion homeostasis,” “excitatory synapse”). These findings, together with downregulation of the intracellular calcium regulator *Calb2,* suggest that genes involved in regulating neuronal excitability may be altered during fentanyl withdrawal. Another notable hub gene was *Galr1*, a member of downregulated module P39 that encodes the galanin receptor. Galanin has been implicated in substance use disorders through interactions with the dopamine and opioid systems in the VTA (Picciotto, 2008). While galanin has been shown to modulate behavioral signs of morphine withdrawal through signaling in the locus coeruleus (Zachariou et al., 2003), our findings identify *Galr1* as a candidate through which galanin signaling may also contribute to fentanyl withdrawal-related adaptations in the VTA.

Consistent with the overall concordance in differential expression between VTA-NAc and VTA-Amy, these projections also showed similarities in the biological processes associated with fentanyl-responsive genes and in the functions of several hub genes. Both projections contained modules enriched for GO terms relating to cellular metabolism and protein processing. In addition, VTA-NAc and VTA-Amy included hub genes with related functions, such as DNA repair (*Zswim7*, N40; *Elof1*, A26), nuclear mRNA transport (*Gle1,* N55; *Nxf1*, A26), and protein ubiquitination (*Fbxw5*, N40; *Usp43*, A26; *Marchf7*, A21). Interestingly, VTA-Amy also showed enrichment for processes related to monoamine signaling that were not prominent in the other projection populations. Upregulated VTA-Amy modules included GO terms associated with monoamine & catecholamine transport, and the top upregulated module A23 contained hub genes encoding DOPA decarboxylase (*Ddc*), vesicular monoamine transporter 2 (*Slc18a2*), and 5-HT receptor 1A (*Htr1a*). Regulatory enrichment analysis further identified *Nhlh1* and *Neurod1* as candidate transcriptional regulators associated with VTA-Amy modules. Both NHLH1 (Gomez Ramos et al., 2023) and NEUROD1 (Rivetti Di Val Cervo et al., 2017) have established roles in dopamine neuron differentiation, raising the possibility that these regulators contribute to the expression of monoamine-related genes during fentanyl withdrawal. Together, these findings suggest that monoaminergic processes may be particularly affected in VTA-Amy neurons during withdrawal and identify dopamine-related signaling in the amygdala as an area for further investigation.

### Transcription-related adaptations are prominent in VTA-NAc during fentanyl withdrawal

A distinct feature of VTA-NAc fentanyl-responsive modules was enrichment for transcription-related functions. For example, the top upregulated VTA-NAc module included the GO term “positive regulation of DNA-templated transcription,” and contained several hub genes encoding transcription factors or transcriptional regulators. This pattern was less prominent in VTA-PFC and VTA-Amy modules, suggesting that genes involved in transcriptional control may be particularly engaged in VTA-NAc neurons during fentanyl withdrawal. Consistent with this possibility, regulatory enrichment analysis identified several transcription factors associated with multiple VTA-NAc fentanyl modules. Among the candidates, *Jun*, an upregulated VTA-NAc DEG, was predicted to regulate 8 VTA-NAc fentanyl modules, while *Gabpb1*, a downregulated VTA-NAc DEG, was predicted to regulate 7 VTA-NAc fentanyl modules. By comparison, transcription factors identified among VTA-PFC or VTA-Amy DEGs were generally associated with only 1-3 fentanyl modules each. These findings raise the possibility that fentanyl withdrawal engages broader transcriptional regulatory networks in VTA neurons projecting to NAc than those projecting to PFC or Amy.

Several transcription factor-encoding genes identified in our dataset have also been implicated previously in substance use-related neuroadaptations, including *Nfkb2*, *Egr1*, and *Jun* (Teague and Nestler, 2022). *Nfkb2* was upregulated in VTA-PFC following fentanyl exposure. While NFkB is best known for its role in inflammatory signaling, NFkB signaling has been shown to promote expression of opioid receptors and peptides that influence drug-associated behaviors (Nennig and Schank, 2017). In VTA-NAc, *Egr1* was upregulated following fentanyl exposure. EGR1 has been implicated in heroin-induced transcriptional adaptations across multiple brain regions (Browne et al., 2023) and is increased in the VTA during morphine withdrawal (García-Pérez et al., 2019). *Jun* was also upregulated in VTA-NAc and was among the candidate regulators associated with the largest number of fentanyl-responsive VTA-NAc modules. JUN can heterodimerize with FOS family proteins to form activator protein-1 (AP-1), a transcription factor complex with well-established roles in neuronal response to drugs of abuse (Nestler, 2008; Browne et al., 2020). Although *Fos* and *Fosb* were not significantly altered by fentanyl in our dataset, both were predicted to regulate 7-8 VTA-NAc fentanyl modules each. Similarly, *Creb1* was not differentially expressed but was predicted to regulate of 5 VTA-NAc fentanyl modules, consistent with the broader involvement of CREB-related signaling in addiction-related transcriptional plasticity (Olson, 2005; Teague and Nestler, 2022). Given that several of these canonical addiction-related transcription factors were preferentially associated with VTA-NAc, it is tempting to speculate that transcriptional adaptations historically attributed to the VTA as a whole may be disproportionately driven by NAc-projecting VTA neurons. Our data do not directly test this possibility, but they provide a projection-specific framework for revisiting established VTA mechanisms of drug-induced plasticity. Future studies will be necessary to determine whether these candidate regulators causally shape transcriptional and behavioral adaptations during opioid withdrawal.

### Conclusions

Together these findings demonstrate that acute spontaneous fentanyl withdrawal produces distinct transcriptional adaptations across VTA neurons that project to either PFC, NAc, and Amy. While VTA-NAc and VTA-Amy showed broadly similar patterns of translatome reprogramming, VTA-PFC exhibited more distinct changes in genes associated with neuronal plasticity, excitability, and neuropeptide signaling. Regulatory enrichment analyses further identified candidate transcriptional regulators associated with projection-specific fentanyl-responsive gene modules, with particularly broad predicted regulatory relationships in VTA-NAc. These findings highlight the importance of projection identity in shaping VTA molecular responses to opioid exposure and identify several promising candidate genes and regulatory mechanisms for future circuit-level studies.

## Supporting information

Supplemental_cluster

Supplemental_PFC

Supplemental_NAc

Supplemental_Amy

Supplemental_input

Supplemental_regulon

## Acknowledgements

The authors thank the Penn State College of Medicine Genome Sciences Core for assistance with RNAseq (RRID:SCR_021123).

## Author Contributions

AM and MEF designed the experiments. AM, HS, LDF, AB, AEO and MEF performed behavior experiments. MEF performed RNA isolation. AM, HS, MEH, and MEF performed data analysis. AM, MEH, and MEF wrote the manuscript and prepared the figures with input from all authors.

## Data and code availability

RNAseq data have been deposited at GEO and are publicly available as of the date of publication (GSE313440). Differential expression, WGCNA, and all code are available at github.com/mfoxlab.

