## Supplemental for "Projection-defined ventral tegmental area neurons exhibit distinct molecular adaptations during acute fentanyl withdrawal"

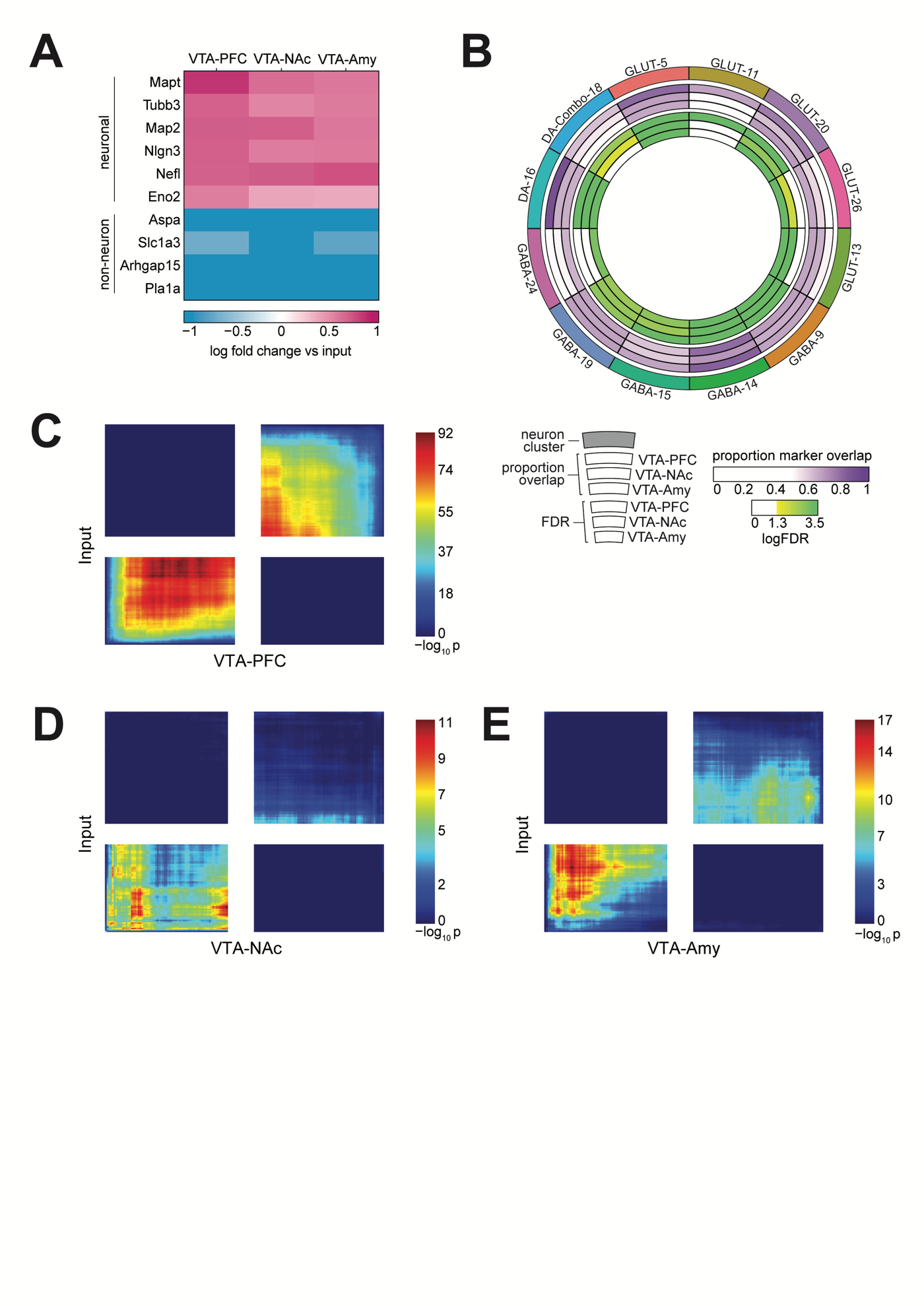
**Supplemental Figure 1. (A)** Heatmap showing enrichment for neuron subtype markers and depletion of non-neuronal markers in VTA-PFC, VTA-NAc, and VTA-Amy saline samples relative to saline input samples. **(B)** Circos plot showing overlap between projection-enriched differentially expressed genes (saline pull-down vs saline input) and marker genes from previously identified mouse VTA neuron clusters. The outer ring indicates VTA neuron clusters classified by dopamine (DA), glutamate (GLUT), or GABA marker expression. For each projection, proportion overlap represents the fraction of differentially expressed genes overlapping any VTA cluster marker. Corresponding Fisher’s exact test logFDR are also shown internally for VTA-PFC, VTA-NAc, and VTA-Amy, respectively. **(C)** RRHO plots showing concordance of fentanyl-induced DEGs between input and VTA-PFC, **(D)** between input and VTA-NAc, **(E)** and between input and VTA-Amy.

**Supplemental File Table of Contents**

| File Name | Sheet Name | Description |
| --- | --- | --- |
|  | [Gene_summary_legends](applewebdata://5152265B-833F-4A2D-B365-4F792B7B81FB#Gene_summary_legends!A1) | Explanation of column names in differential expression summary sheets |
| Supplemental_File_NAc.xlsx | [VTA-NAc_gene_summary](applewebdata://5152265B-833F-4A2D-B365-4F792B7B81FB#'VTA-NAc_gene_summary'!A1) | differential expression analysis, module assignment, and correlation with module eigengene for individual genes in VTA-NAc WGCNA |
| Supplemental_File_PFC.xlsx | [VTA-PFC_gene_summary](applewebdata://5152265B-833F-4A2D-B365-4F792B7B81FB#'VTA-PFC_gene_summary'!A1) | differential expression analysis, module assignment, and correlation with module eigengene for for individual genes in VTA-PFC WGCNA |
| Supplemental_File_AMY.xlsx | [VTA-AMY gene summary](applewebdata://5152265B-833F-4A2D-B365-4F792B7B81FB#'VTA-Amy_gene_summary'!A1) | differential expression analysis, module assignment, and correlation with module eigengene for for individual genes in VTA-Amy WGCNA |
| Supplenmental_File_INPUT.xlsx | [Input_gene_summary](applewebdata://5152265B-833F-4A2D-B365-4F792B7B81FB#Input_gene_summary!A1) | differential expression analysis for individual genes in input fraction |
| Supplemental_File_GO.xlsx | [Gprofiler_column_legends](applewebdata://5152265B-833F-4A2D-B365-4F792B7B81FB#Gprofiler_column_legends!A1) | Explanation of column names in gprofiler gene ontology analysis sheets |
|  | [VTA-NAC_GO_concat](applewebdata://5152265B-833F-4A2D-B365-4F792B7B81FB#'VTA-NAc_GO_concat'!A1) | Gene Ontology Analysis of genes in each fentanyl-associated VTA-NAc module, obtained with gprofiler2 |
|  | [VTA-PFC_GO-concat](applewebdata://5152265B-833F-4A2D-B365-4F792B7B81FB#'VTA-PFC_GO_concat'!A1) | Gene Ontology Analysis of genes in each fentanyl-associated VTA-PFC module, obtained with gprofiler2 |
|  | [VTA-AMY_GO-concat](applewebdata://5152265B-833F-4A2D-B365-4F792B7B81FB#'VTA-Amy_GO_concat'!A1) | Gene Ontology Analysis of genes in each fentanyl-associated VTA-AMY module, obtained with gprofiler2 |
| Supplemental_file_Regulon.xlsx | [iRegulon_legends](applewebdata://5152265B-833F-4A2D-B365-4F792B7B81FB#iRegulon_legends!A1) | Parameters for iRegulon analysis and explanation of column names in regulator summary sheets |
|  | [VTA-NAc-module-regulators](applewebdata://5152265B-833F-4A2D-B365-4F792B7B81FB#'VTA-NAc-module-regulators'!A1) | Predicted transcription factors for protein coding genes from selected VTA-NAc modules, obtained with iRegulon |
|  |  | [VTA-NAc_Regulon_N34](applewebdata://5152265B-833F-4A2D-B365-4F792B7B81FB#'VTA-NAc_Regulon_N34'!A1) |
|  |  | [VTA-NAc_Regulon_N40](applewebdata://5152265B-833F-4A2D-B365-4F792B7B81FB#'VTA-NAc_Regulon_N40'!A1) |
|  |  | [VTA-NAc_Regulon_N46](applewebdata://5152265B-833F-4A2D-B365-4F792B7B81FB#'VTA-NAc_Regulon_N46'!A1) |
|  |  | [VTA-NAc_Regulon_N55](applewebdata://5152265B-833F-4A2D-B365-4F792B7B81FB#'VTA-NAc_Regulon_N55'!A1) |
|  | [VTA-PFC-module-regulators](applewebdata://5152265B-833F-4A2D-B365-4F792B7B81FB#'VTA-PFC-module-regulators'!A1) | Predicted transcription factors for protein coding genes from selected VTA-PFC modules, obtained with iRegulon |
|  |  | [VTA-PFC_Regulon_P5](applewebdata://5152265B-833F-4A2D-B365-4F792B7B81FB#'VTA-PFC_Regulon_P5'!A1) |
|  |  | [VTA-PFC_Regulon_P9](applewebdata://5152265B-833F-4A2D-B365-4F792B7B81FB#'VTA-PFC_Regulon_P9'!A1) |
|  |  | [VTA-PFC_Regulon_P10](applewebdata://5152265B-833F-4A2D-B365-4F792B7B81FB#'VTA-PFC_Regulon_P10'!A1) |
|  |  | [VTA-PFC_Regulon_P39](applewebdata://5152265B-833F-4A2D-B365-4F792B7B81FB#'VTA-PFC_Regulon_P39'!A1) |
|  | VTA-Amy-module_regulators | Predicted transcription factors for protein coding genes from selected VTA-Amy modules, obtained with iRegulon |
|  |  | [VTA-Amy_Regulon_A19](applewebdata://5152265B-833F-4A2D-B365-4F792B7B81FB#'VTA-Amy_Regulon_A19'!A1) |
|  |  | [VTA-Amy_Regulon_A21](applewebdata://5152265B-833F-4A2D-B365-4F792B7B81FB#'VTA-Amy_Regulon_A21'!A1) |
|  |  | [VTA-Amy_Regulon_A23](applewebdata://5152265B-833F-4A2D-B365-4F792B7B81FB#RANGE!A1) |
|  |  | [VTA-Amy_Regulon_A26](applewebdata://5152265B-833F-4A2D-B365-4F792B7B81FB#RANGE!A1) |
|  | [TF intersections column legends](applewebdata://5152265B-833F-4A2D-B365-4F792B7B81FB#RANGE!A1) | Explanation of column names in transcription factor (TF) intersections sheet |
|  | [TF intersections](applewebdata://5152265B-833F-4A2D-B365-4F792B7B81FB#RANGE!A1) | Compiled differential expression information for predicted TFs of fentanyl-associated modules across all VTA projections |
| Supplemental_File_Clustermarkers.xlsx | [Cluster_markers_column_legends](applewebdata://5152265B-833F-4A2D-B365-4F792B7B81FB#cluster_markers_column_legends!A1) | Explanation of column names in cluster marker sumary sheet |
|  | [Cluster_markers](applewebdata://5152265B-833F-4A2D-B365-4F792B7B81FB#cluster_markers!A1) | Differentially expressed genes overlapping with cluster markers from snRNAseq from DOI:10.1111/adb.13403, as determined using the MeanRatio method from DeconvoBuddies (DOI: 10.1186/s13059-025-03552-3) |
